# Bats adjust temporal features of echolocation calls but not those of communication calls in response to traffic noise

**DOI:** 10.1101/354845

**Authors:** Shengjing Song, Aiqing Lin, Tinglei Jiang, Xin Zhao, Walter Metzner, Jiang Feng

## Abstract

**Summary statement:** This study reveals the impact of anthropogenic noise on spectrally distinct vocalizations and the limitations of the acoustic masking hypothesis to explain the vocal response of bats to chronic noise.

**Abstract:** The acoustic masking hypothesis states that auditory masking may occur if the target sound and interfering sounds overlap spectrally, and it suggests that animals exposed to noise will modify their acoustic signals to increase signal detectability. However, it is unclear if animals will put more effort into changing their signals that spectrally overlap more with the interfering sounds than when the signals overlap less. We examined the dynamic changes in the temporal features of echolocation and communication vocalizations of the Asian particolored bat (*Vespertilio sinensis*) when exposed to traffic noise. We hypothesized that traffic noise has a greater impact on communication vocalizations than on echolocation vocalizations and predicted that communication vocalization change would be greater than echolocation. The bats started to adjust echolocation vocalizations on the fourth day of noise exposure, including an increased number of call sequences, decreased number of calls, and vocal rate within a call sequence. However, there was little change in the duration of the call sequence. In contrast, these communication vocalization features were not significantly adjusted under noise conditions. These findings suggest that the degree of spectral overlap between noise and animal acoustic signals does not predict the level of temporal vocal response to the noise.

## INTRODUCTION

Anthropogenic noise disturbs acoustic communication, prey detection, and predator avoidance in many animals (e.g. Senzaki et al., 2016; Siemers and Schaub, 2011; Templeton et al., 2016; Brumm and Slabbekoorn, 2005; Francis and Barber, 2013; Kight and Swaddle, 2011; Shannon et al., 2016), and animal response to noise is an important topic in ecology and conservation. Many studies have demonstrated that animals can adjust their vocalizations in response to ambient noise (e.g. Brumm, 2004; Duarte et al., 2018; Hage et al., 2013; Lowry et al., 2013; Parks et al., 2011; Parris et al., 2009). Some groups, such as birds and bats, can have multiple types of vocalizations that are spectrally distinct (e.g. Catchpole and Slater, 1995; Kanwal et al., 1994; Kroodsma, 2004; Lin et al., 2016). However, little is known about the differences in the noise-induced acoustic adjustments among vocalization types.

Noise can negatively impact animal vocalization by acoustic masking, attention reduction, and increased stress (Francis and Barber, 2013; Kight and Swaddle, 2011). Among these impacts, acoustic masking is especially problematic and has been used to explain the vocal response to noise for many species (e.g. Brumm et al., 2004; Corcoran and Conner, 2014; Francis et al., 2011; Gerhardt and Klump, 1988). The acoustic masking hypothesis states that auditory masking may occur if the target sound and interfering sounds overlap in frequency (Luo et al., 2015b). Noise can reduce the transmission effectivity and detectability of acoustic signals (Brumm and Slabbekoorn, 2005). To reduce the negative effect of acoustic masking, animals may adjust the spectral and/or temporal features of their vocalizations by increasing amplitude, frequency, bandwidth and duration of vocal elements (Hage et al., 2013; Parks et al., 2007; Rios-Chelen et al., 2012; Siegert et al., 2013), and increasing the number of vocal elements and repetition rate (Bittencourt et al., 2017; Brumm et al., 2004; Caldart et al., 2016; Luther and Gentry, 2013; Nelson et al., 2017; Roy et al., 2011). Since masking is most effective when the masking sound spectrally overlaps with the target sound, it is expected that noise would have more negative impacts on the vocalizations that spectrally overlap more with the interfering noise and that animals under noisy conditions would make a greater effort to change their vocalizations than those that spectrally overlapped less with the noise.

Bats are nocturnal flying mammals that rely mainly on acoustic signals for environmental perception and social communication. Bats that echolocate not only produce echolocation signals for navigation, orientation, and prey detection (Jones and Teeling, 2006; Moss et al., 2011), but they also have rich vocal repertoires for individual recognition, mate attraction, predator avoidance, and territorial defense (Bohn et al., 2008; Gadziola et al., 2012). Echolocation calls are spectrally distinct from communication calls. Echolocation calls are relatively simple in spectrum with most energy concentrated at frequencies ranging from 20 to 150 kHz (Jones and Teeling, 2006). In contrast, communication calls are often spectrally complex with most energy concentrated at frequencies below 25 kHz (Bohn et al., 2008; Gadziola et al., 2012; Kanwal et al., 1994; Lin et al., 2016; Ma et al., 2006).

Some bats inhabit or forage near man-made structures, such as bridges, buildings, mines, and highways, which are noisy (Kunz and Fenton, 2006). Bats are among the vertebrates that suffer the most from noise pollution. Noise can decrease the foraging activity and efficiency of bats (Bunkley and Barber, 2015; Bunkley et al., 2015; Luo et al., 2015b; Schaub et al., 2008; Siemers and Schaub, 2011). In response to noise, bats can adjust the acoustic characteristics of their echolocation calls, such as amplitude, frequency, and duration (e.g. Bunkley et al., 2015; Hage et al., 2013; Hage and Metzner, 2013; Luo et al., 2015a; Luo and Moss, 2017). However, little is known about the impact of noise on bat communication vocalizations. Moreover, the current knowledge on bat vocal adjustment to noise is based on short-term noise exposure, usually for only several minutes or less. It is unknown how bats might change their vocalizations when exposed to chronic noise lasting several days or longer.

We studied how bats might change the temporal features of echolocation and communication vocalizations when they are exposed to chronic noise. Asian particolored bats (*Vespertilio sinensis*) were collected from a colony that roosted beneath a traffic bridge and were exposed to the loud noises and vibrations associated with the passing vehicles. The bats in this bridge colony frequently emitted echolocation and communication calls with the dominant frequency of echolocation calls around 34 kHz and the majority of communication calls ranging from 9-21 kHz (Fig. 1A, B). Most of the traffic noise energy has a frequency range of 3-15 kHz (Fig. 1E). We tested the hypothesis that traffic noise had a greater negative impact on bat communication vocalizations than echolocation vocalizations. We predicted that, in response to traffic noise, bats would change more in the temporal features of vocalizations for communication than for echolocation. We focused on the temporal rather than the spectral features of the vocalizations for the following reasons: first, the impact of noise on the spectral parameters of vocalizations has been described in bats but the impact on temporal features is unclear. Therefore, temporal features were the focus of the study. Second, the spectral features of vocalizations of freely housed bats are difficult to qualify because these features are easily affected by distance and the direction of recording.

**Figure 1.**
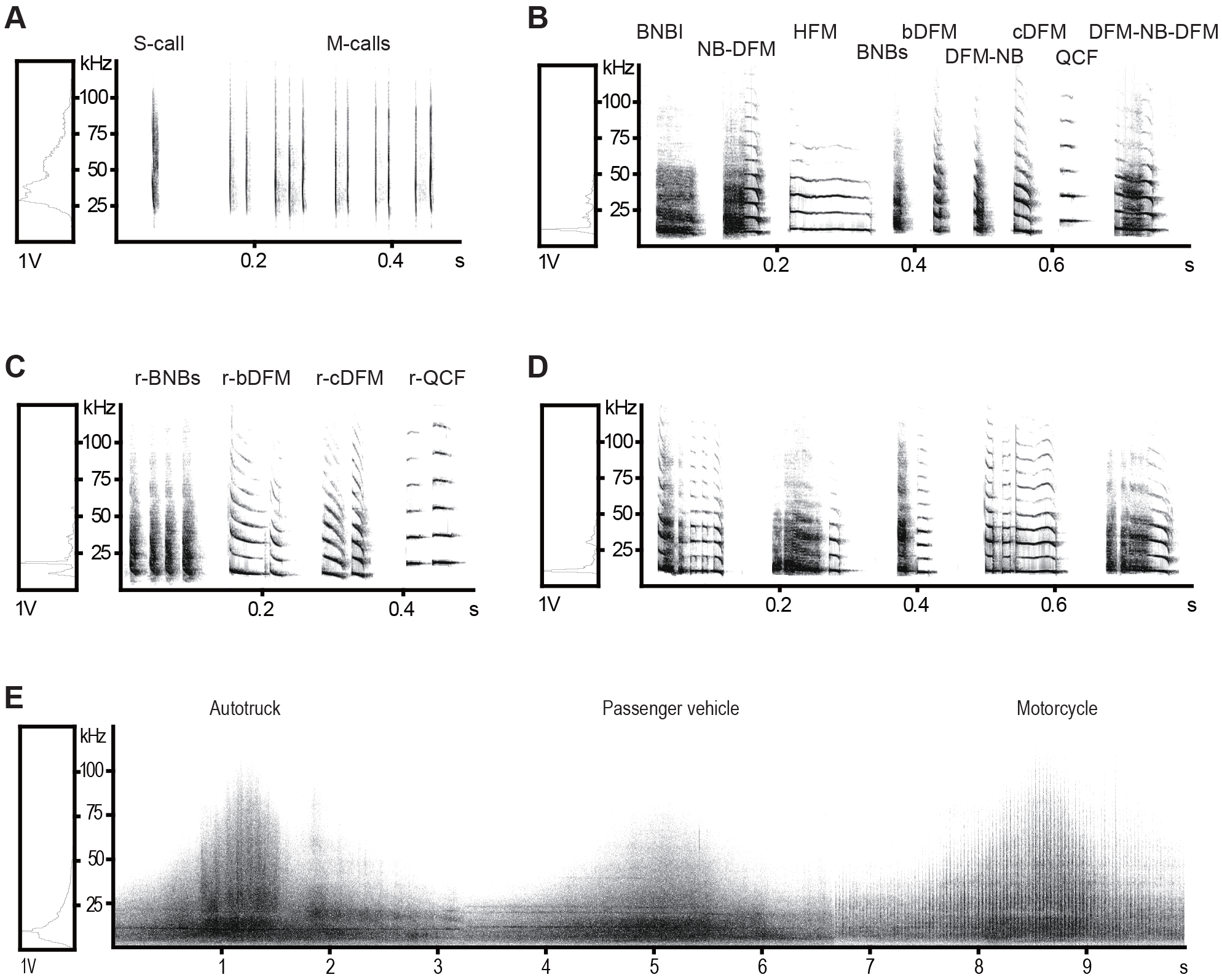
Spectrograms of vocalizations of *Vespertilio sinensis* and traffic noise. (A) An example of echolocation single-call sequence and multiple-calls sequence. (B) Nine distinct syllabic types of communication vocalizations (see Table S1 for detailed abbreviation adopted for each syllable). (C) The most frequently observed types of communication single-syllable call sequences (SS-call sequences). (D) An example of communication composite-syllable call sequence (CS-call sequence). (E) Traffic noise produced by the most common types of vehicles passing over the bridge where the *V. sinensis* roost.

## MATERIALS AND METHODS

### Location and animals

Twenty-six non-pregnant female adult *V. sinensis* were collected from a large colony inhabiting a traffic bridge in Harerbin City, China. The bat colony roosts under the bridge from June to October each year and has more than 10,000 individuals. The traffic volume on the bridge is high with more than 500 vehicles per hour during the daytime (8:30–18:30). Trucks represent more than 25% of the vehicles crossing the bridge.

The collected bats were housed at Northeast Normal University, Changchun, China. The bats were divided into two groups comprised of individuals with similar body size (forearm length: noise-exposure group 49.33 ± 1.39 mm; control group 50.45 ± 1.93 mm). The two groups were separately housed in cages (100 cm long × 60 cm wide × 80 cm high) in isolated rooms under the same conditions of temperature and illumination (temperature: 22−24 °C; relative humidity: 50%−60%; light cycle 12-h light/12-h dark, 07: 00−19: 00 for light conditions; 4.80 m long × 2 m wide × 2.50 m high). They were given ad libitum access to water and mealworms, and their diet was enriched with vitamin and mineral supplements.

### Noise recording and generation of playback files

We recorded traffic noise on the bridge for acoustic playback. To obtain high-quality traffic noise, an ultrasonic microphone (UltraSoundGate CM16/CMPA, Avisoft Bioacoustics, Berlin, Germany) was placed at a distance of about 5 m from the nearest passing vehicles. The microphone was connected to an ultrasound recording interface (UltraSoundGate 116H, Avisoft Bioacoustics), with a sample rate of 375 kHz at 16 bits/sample. Traffic noise was recorded from 08:30 to 18:30 h with a total of 10 days of recording. The recorded files were automatically saved with a one-minute duration of each file. Each file was high-pass filtered at 1 kHz using Avisoft-SASLab Pro 5.2.

We measured the sound level of traffic noise in the location inhabited by the bats as a reference of amplitude for noise playback. A calibrated microphone (46BF, G.R.A.S. Sound & Vibration, Denmark) was positioned about 50 cm from the bats in the crevices under the bridge. The microphone was connected to an ultrasound recording interface (UltraSoundGate 116Hm, Avisoft Bioacoustics), with a sample rate of 375 kHz at 16 bits/sample. A 30-minute recording (10:00–10:30 h) showed that the traffic noise was about 63 (range = 60–65) dB SPL (sound pressure level re. 20 μPa). The noise the bats experienced was louder than this level since the noise caused by vehicles can transmit via the bridge structure in addition to the air.

Fifty silence files, each one-minute long, were created using Avisoft-SASLab Pro 5.2, with a sample rate of 375 kHz at 16 bits/sample. These silence files did not have any acoustic signals and were broadcast to the bats of control group.

### Acoustic playback and sound recording

In the first 14 d, the bats of both groups were maintained without any sound recording or playback. We then recorded the vocalizations of the bats to determine if there were any differences in the vocalizations of the two groups with and without noise exposure. We recorded bat vocalizations from 08:30 to 18:30 h each day and achieved a total of 5 d of recordings. Vocalizations were recorded using an ultrasonic microphone (UltraSoundGate CM16/CMPA) and interface (UltraSoundGate 116H), with a sample rate of 375 kHz at 16 bits per sample. Each microphone was placed at a distance of one meter from the middle of the cage.

We then broadcast files of traffic noise to the bats of the noise-exposure group and silence files to the control group. Playback files were played by a loudspeaker (Ultrasonic Dynamic Speaker, Vifa, Avisoft Bioacoustics), which was positioned at a distance of three meters from the cage. Each speaker was connected to an ultrasound playback interface (UltraSoundGate player 116, Avisoft Bioacoustics). Traffic noise were played with an amplitude of around 63 dB (60–65 dB) at the sites where the bats were hanging. A microphone (UltraSoundGate CM16/CMPA), connected to a recording interface (Avisoft UltraSoundGate 116H), was positioned at a distance of one meter from the cage to record the vocalizations of the bats. The direction of the microphone was consistent with that of the loudspeaker. Bat vocalizations were recorded with a sample rate of 375 kHz at 16 bits per sample. Traffic noise and silence files were respectively broadcast to the bats of noise exposure group and control group during 08:30–18:30 h each day for a total of 10 d of playback. The recorded noise files were sorted randomly by day and played to the bats in the noise-exposure group. The 50 silence files were randomly broadcast to the control group bats.

### Data analysis

The vocalizations of the bats consist of call sequences. A “call” is defined as a discrete vocalization, which is surrounded by periods of substantial silence when the call is emitted by itself. It is regarded as the smallest, acoustically independent, unit of a vocalization. A “call” here is the same as the “pulse” of echolocation vocalizations and the “syllable” of communication vocalizations (Kanwal et al., 1994). A call sequence consists of a single call (S-call sequence) or multiple calls (M-calls sequence). We measured the inter-call interval (duration between the end of a call and the onset of next one) to establish the boundary of the call sequence. Successive two calls were randomly selected from the data base of vocalizations recorded and 1100 call intervals were measured respectively for echolocation and communication vocalizations of each bat group. *V. sinensis* often emit two echolocation calls each time, which results in a trimodal distribution of call intervals of echolocation vocalizations (Fig. S1A). We set the second trough as a boundary of call sequence so that two successive calls were classified into two different call sequences when their interval was greater than 250 ms (Fig. S1A) or into the same call sequence if the interval shorter than 250 ms. Similarly, the inter-call interval of 40 ms was set as a call sequence boundary for communication vocalizations (Fig. S1B).

A total of 92,296 call sequences of echolocation and communication vocalizations were analyzed for the noise-exposure group and control group. We counted the daily number of echolocation call sequences of each bat group: the number of S-call sequences, the number of M-calls sequences, and the total number of call sequences (including both S-call sequences and M-calls sequences). In addition, we analyzed the following parameters: the duration of an S-call sequence, and the duration, number of calls, and vocal rate within an M-calls sequence. The vocal rate is the number of calls divided by the duration of the call sequence. A daily mean value was calculated for each parameter.

*V. sinensis* produce multiple types of communication syllables (calls) (Fig. 1B-D). We classified syllabic types following the methods reported in Kanwal et al. (1994). We then classified the communication call sequences into three categories: single-syllable call sequence (SS-call sequence: a call sequence consists of a single syllable), repeat-syllable call sequence (RS-call sequence: a call sequence consists of repeats of the same syllables), composite-syllable call sequence (CS-call sequence: a call sequence consists of multiple types of syllables). The numbers of communication call sequences per day in each bat group were analyzed as follows: the number of SS-call sequences, the number of RS-call sequences, the number of CS-call sequences, and the total number of call sequences (including SS-call sequences, RS-call sequences, and CS-call sequences). We analyzed the duration of each SS-call sequence, the duration, number of calls, vocal rate within each RS-call sequence and each CS-call sequence. A daily mean value was calculated for each parameter. For SS-call sequences and RS-call sequences, only the call sequences consisting of the most frequently observed syllables were selected for statistical analysis.

All of the acoustic parameters were measured from spectrograms, with the Hamming window FFT = 512, overlap = 93.73%, using Avisoft-SASLab Pro v5.2. The normality of data was tested using Kolmogorov-Smirnov test. The differences in daily mean values of each parameter between bats of noise-exposure and control groups were tested using an Independent sample *T-test* or the Mann-Whitney *U-test.* All of the statistical analyses were performed by SPSS 22 (IBM). To minimize bias, a blinded method was used so that the person who analyzed the acoustic data had no knowledge of the bat treatment.

## RESULTS

Vocal activities were not significantly different between bats of the noise exposure group and control group during the first 5 d before playback. There was no significant difference in the daily total number of echolocation call sequences (noise exposure group: 1816 ± 42.61, control group: 1771.20 ± 256.73; Independent samples T test, *P* > 0.05; Fig. 2A), the daily number of S-call sequences (noise-exposure group: 577.20 ± 26.78; control group: 563.00 ± 4.00; Independent samples T test, *P* > 0.05; Fig. 2B), and the ratio of M-calls sequences to the total number of echolocation call sequences per day (noise-exposure group: 68.24 ± 6.58%; control group: 67.81 ± 6.58%; Independent samples T test, *P* > 0.05). The daily total number of communication call sequences (noise exposure group: 1135.60 ± 417.09; control group: 1097.40 ± 233.47), the daily number of SS-call sequences (noise exposure group: 82.49 ± 3.58%; control group: 76.71 ± 6.28%), the daily number of RS-call sequences (noise exposure group: 5.04 ± 0.63%; control group: 4.88 ± 1.98%), and the ratio of the number of CS-call sequences to the total number of call sequences per day (noise exposure group: 12.47 ± 3.83%; control group: 14.40 ± 7.74%) were not significantly different between the two groups before playback (Independent samples T test, all *P* > 0.05).

**Figure 2.**
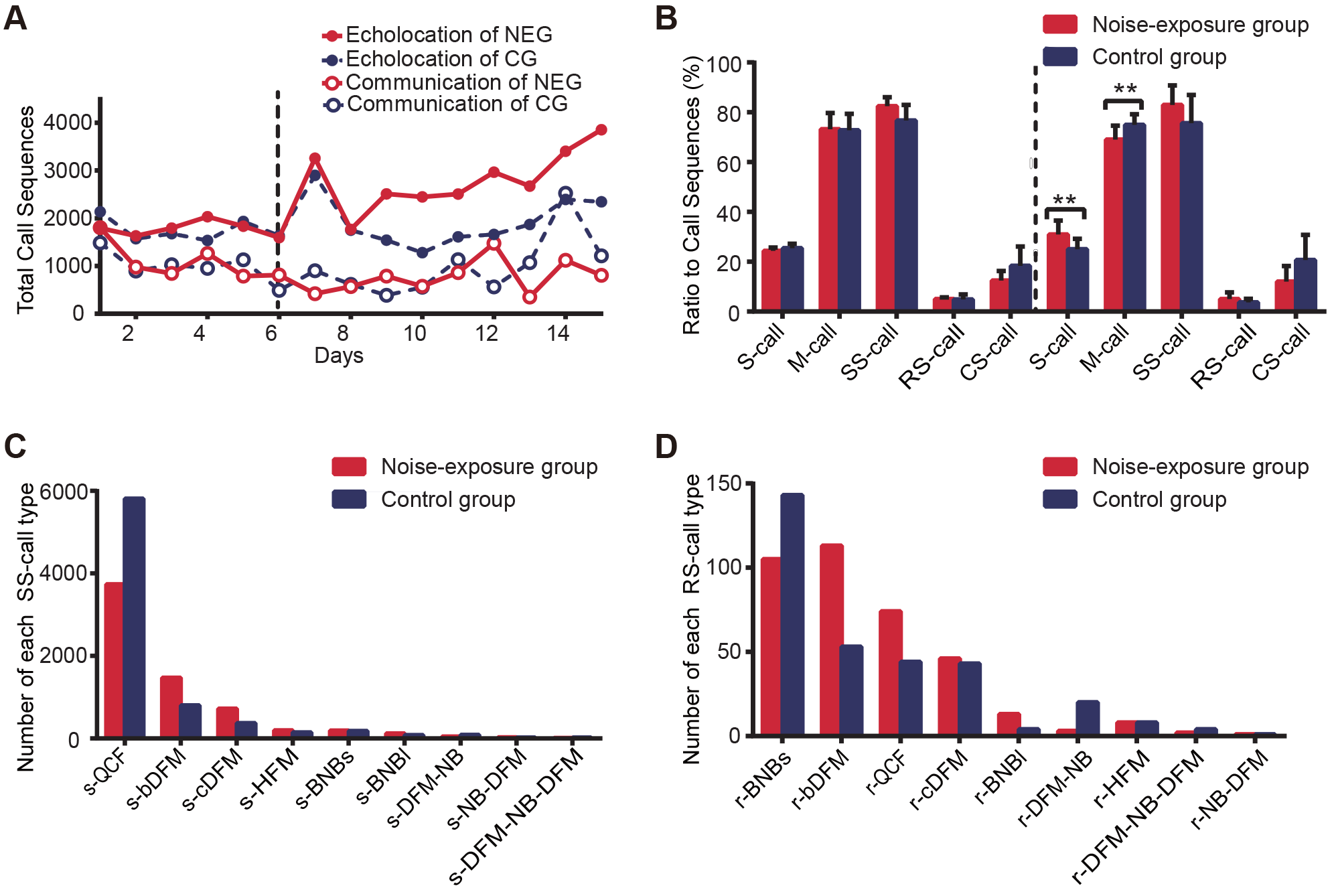
Temporal features of echolocation and communication vocalizations in *Vespertilio sinensis* in the noise-exposure group (NEG) and control group (CG) before and after playback. (A) The daily total number of call sequences of echolocation (filled circle) and communication (open circle) vocalizations in bats in the noise-exposure group (red) and control group (blue). (B) The ratio of the number of single-call sequence (S-call) or multiple-calls sequence (M-calls) to the total number of echolocation call sequences, and the ratio of the number of single-syllable call sequences (SS-call), repeat-syllable call sequence (RS-call), and composite-syllable call sequence (CS-call) to the total number of communication call sequences. (C) and (D) The average daily total number of each type of communication single-call sequences in noise-exposure group (red) and control group (blue). The vertical dotted line in (A) and (B) indicates the results obtained before (left) and after (right) playback. Asterisk indicates a statistically significant difference (* P<0.05, ** P<0.01). See Table S1 for detailed abbreviation adopted for each syllable.

### Numbers of Call Sequences

The bats under noise exposure were more active in echolocation vocalization compared with those under silence playback. The average daily total number of echolocation call sequences of the noise-exposure group was 29.61% greater than that of the control group during the whole period of playback (noise exposure group: 2697.90 ± 702.89; control group: 1899.00 ± 15.00; Independent samples T test, *F* = 0.91, *P* = 0.001; Fig. 2A). On the first 3 d of playback, the differences in the total number of echolocation call sequences per day between the two groups of bats ranged from 0.80%-12.31%. After 3 d, the differences increased and ranged from 42.19% to 91.25% (Fig. 2A). Bats exposed to noise emitted more S-call sequences but fewer M-calls sequences per day, with an increased ratio of the number of S-call sequences to the total number of call sequences (Independent samples T test, *F* = 2.85, *P* = 0.015; Fig. 2B).

There was no significant difference in the communicative vocal activity between the two groups during the period of playback. The daily total number of communication call sequences (Independent samples T test, *F* = 1.57, *P* = 0.464; Figure. 2A) and the ratio of the number of call sequences of each call type to the total number of call sequences were not significantly different between the bats under noise exposure and silence conditions (Independent samples T test: SS-call sequence: *F* = 4.28, *P* = 0.110; RS-call sequence: *F* = 3.89, *P* = 0.200; CS-call sequence: Mann-Whitney *U*-test, *Z* = 1.97, *P* = 0.052; Figure. 2B). SS-call sequences accounted for about 80% of the daily total number of communication call sequences (noise exposure group: 82.96 ± 7.84%; control group: 75.61 ± 11.40%). The call sequences of s-QCF, s-bDFM and s-cDFM were the most frequently observed SS-call sequences, accounting for about 92% of the total number of SS-call sequences (Fig. 2C). The call sequences of r-BNBs, r-bDFM, r-QCF and r-cDFM accounted for about 78% of the number of RS-call sequences (Fig. 2D). These seven most common call sequences as well as the CS-call sequences were selected for further analyses. The abbreviations for each syllable are shown in Table S1.

### Duration of Call Sequences

There was no significant difference in the duration of an echolocation S-call sequence (noise exposure group: 5.14 ± 1.15 ms; control group: 5.59 ± 0.73 ms; Mann-Whitney *U*-test, *Z* = −1.74, *P* = 0.089; Fig. 3A) or an M-calls sequence (noise exposure group: 891.09 ± 96.43 ms; control group: 959.51 ± 121.15 ms; Independent samples T test, *F* = 1.58, *P* = 0.197; Fig. 3B) between the two bat groups. Under the noise condition, only the duration of r-cDFM call sequence was reduced significantly (noise exposure group: 111.88 ± 74.38 ms; control group: 219.77 ± 125.30 ms; Independent samples T test, *F* = 1.94, *P* = 0.031). The bats did not significantly change the duration of a call sequence of all of the remaining communication call types when exposed to noise (all *P* > 0.05; Fig. 3A and 3B).

**Figure 3.**
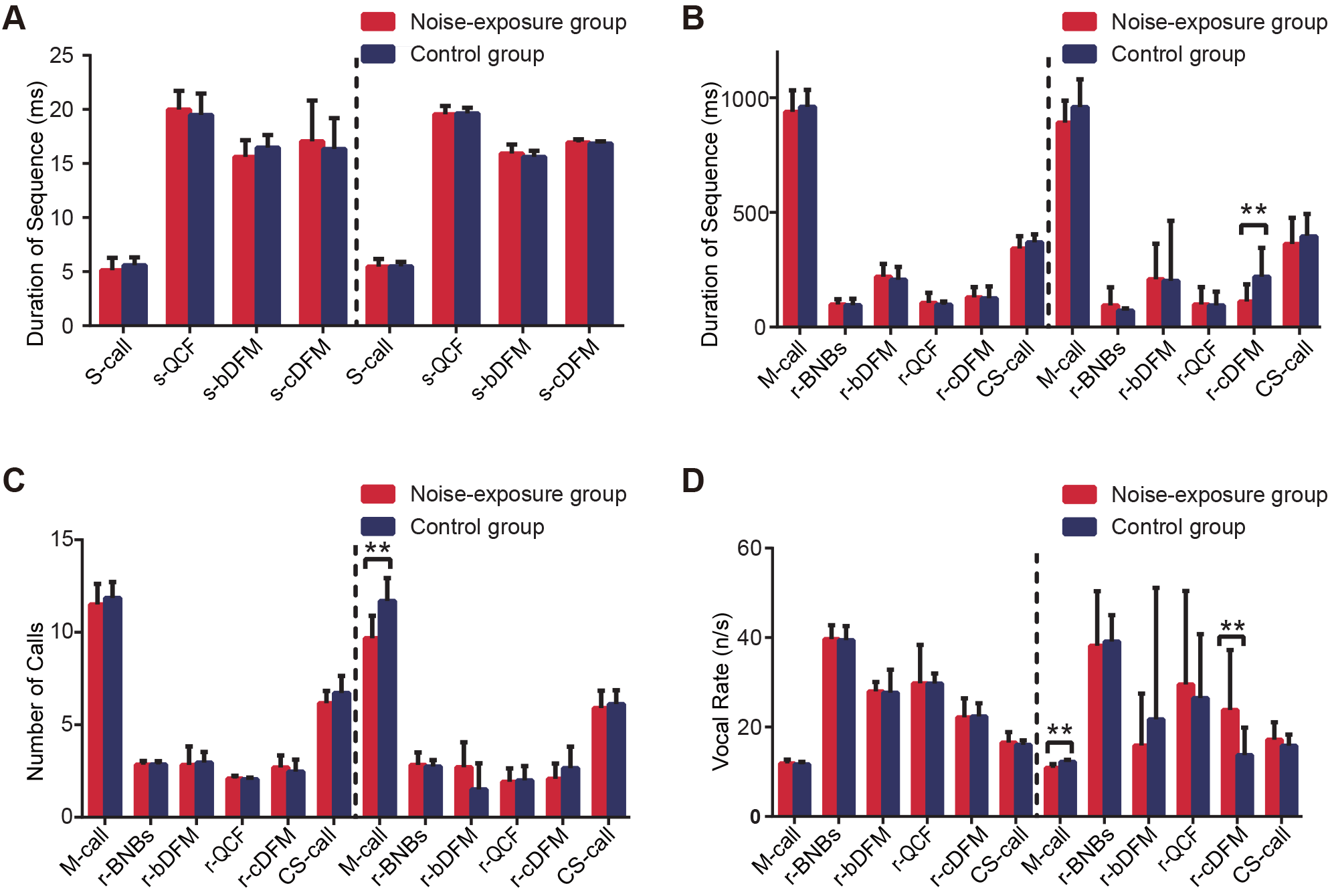
Temporal features of echolocation and communication vocalizations in *Vespertilio sinensis* before and after playback. (A) Duration of echolocation single-call sequence (S-call) and duration of communication single-syllabic call sequence. (B) Duration of echolocation multiple-calls sequence (M-call) and duration of communication repeat-syllable call sequence and composite-syllable call sequence. The number of calls (C) and the vocal rate (D) within an echolocation multiple-calls sequence, within a communication repeat-syllable call sequence, and within a composite-syllable call sequence. The vertical dotted line indicates the results obtained before (left) and after (right) playback. Asterisk indicates a statistically significant difference (* P<0.05, ** P<0.01). See Table S1 for detailed abbreviation adopted for each syllable.

### Number of elements within call sequences

The number of calls within an echolocation M-calls sequence was decreased by 17.2% in the bats of the noise-exposure group and was significantly less than the value of the bats in the control group (Mann-Whitney U-test, *Z* = −2.87, *P* = 0.003; Fig. 3C), resulting in a significantly slower vocal rate within an echolocation call sequence (noise exposure group: 10.88 ± 0.84; control group: 12.22 ± 0.48; Independent samples T test, *F* = 0.56, *P* < 0.001; Fig. 3D). Under noise conditions, the numbers of calls and vocal rate within a call sequence did not change significantly in any communication call type (all *P* > 0.05; Fig. 3C), except for the call sequences of r-cDFM, within which the vocal rate significantly increased by 45% (Mann-Whitney *U*-test, *Z* = −2.16, *P* = 0.031; Fig. 3D).

## DISCUSSION

This study demonstrated that Asian particolored bats responded to traffic noise by changing their echolocation vocalizations. They increased the total number of call sequences, increased the number of single-call sequences, and decreased the number of calls and vocal rate within multiple-calls sequences. In contrast, all but one the most frequently observed communication call types were stable in these features. The total number of communication call sequences as well as the duration, number of calls, and vocal rate within a communication call sequence were not significantly changed under the noise treatment. These results show that the bats adjusted the temporal features of echolocation vocalizations but not those of communication vocalizations when exposed to chronic noise. This result did not support the prediction that bats would change more in the acoustic features of their vocalizations for communication compared to echolocation in response to traffic noise.

### Increased echolocation activity under noise condition

The most obvious vocal response of the bats to traffic noise was an increased number of echolocation call sequences. One possible reason is that the vocal response increases the detectability of echolocation calls. Anthropogenic noise can potentially mask high-frequency vocalizations because energy in the spectral region of a vocal signal also contributes to masking signals in other frequencies, albeit to a lesser extent (Díaz et al., 2011; Parris and Schneider, 2009). Echolocation calls of *V. sinensis* spectrally overlap with the high-frequency parts of the broadcast traffic noise, although the traffic noise has most of its energy in its low frequencies. Traffic noise may reduce the detectability of echolocation signals by acoustic masking. A recent study showed that traffic noise can affect the starting frequency, peak frequency, and ending frequency of echolocation calls in this species under natural conditions (Guo *et al.* 2015). The acoustic masking of echolocation calls by anthropogenic noise in several other bat species has also been suggested (Bunkley et al., 2015).

Many animals can adjust their vocalizations to improve signal detectability under noise conditions by changing spectral (amplitude and frequency) and/or temporal (vocal rate and timing) characteristics (Brumm and Slabbekoorn, 2005). A rise in the number of call sequences in *V. sinensis* may increase the redundancy of the signals that can contribute to improving signal detectability and facilitating signal detection (Brumm and Slabbekoorn, 2005; Luo et al., 2015a; Luther and Gentry, 2013). Similar results have been reported in birds. For example, *Serinus serinus* increase the time spent singing on posts in noisy city environments (Díaz et al., 2011). *V. sinensis* increase the number of single-call sequences but decrease the number of multiple-call sequences, which possibly decreases the complexity of their calls. Acoustic signals with low complexity can increase signal-to-noise ratio and/or have higher transmission efficiency (Dabelsteen et al., 1993; Lohr et al., 2003). *V. sinensis* bats may adjust these acoustic features of echolocation vocalizations to improve the detectability of the signals in response to noise.

Adjustment of echolocation vocalization could also be a stressor response to the traffic noise. Noise exposure may act as a stressor to animals and thereby change physiological characteristics (Kight and Swaddle, 2011; Wright et al., 2007), such as increasing stress hormone levels and inducing immunosuppressive effects that can alter vocal performance (Francis and Barber, 2013; Grunst and Grunst, 2014; Troïanowski et al., 2017). In this study, the bats exposed to noise did not significantly change their echolocation vocalizations until the fourth day of playback. This result differs from other bat studies testing short-term noise conditions where bats altered their vocalizations immediately when exposed to noise (e.g. Hage et al., 2013; Luo et al., 2017a). The different results may have been caused by the different acoustic parameters studied or they may reflect different underlying mechanisms. Improved signal detectability cannot explain the 3-d latency of vocal adjustment in this study. Instead, the vocal-response latency is probably a result of accumulated noise stress and physiological responses. Under stress, behavioral response due to altered physiological processes may be relatively slow and delayed owing to the requirement of accumulating hormones, such as glucocorticoids (Romero and Butler, 2007).

### Different response of echolocation and communication vocalization

The response of echolocation vocalization to noise has been reported in other bat species (Bunkley et al., 2015; Hage et al., 2013; Hage and Metzner, 2013; Luo et al., 2017a; Luo et al., 2017b). However, little is known about the impact of noise on bat communication vocalizations. The extent to which bats will respond to noise differently in echolocation and communication vocalizations is not known. *V. sinensis* bats emit both echolocation and communication calls in noise conditions. It is expected that noise masking of communication vocalizations would be stronger than on echolocation vocalizations and that the bats would alter communication calls more than echolocation calls. In contrast, our results showed that the bats, in response to traffic noise, changed the temporal features of their echolocation calls but not those of communication calls. Although the duration and call rate was adjusted in one type of communication vocalizations, this vocal type accounted for only 1% of the total number of communication call sequences.

One possible reason for this unexpected result is that communication and echolocation vocalizations are different in the acoustic response to noise masking. V *sinensis* can change frequency parameters of echolocation calls under traffic noise conditions (Guo et al., 2015). *V. sinensis* also increases the frequencies and duration of communication calls under noise (Jiang et al., unpublished data). These findings, together with our results, suggest that *V. sinensis* can adjust both spectral and temporal features of their echolocation calls and adjust spectral but not temporal features of communication calls under the influence of noise. The other possible explanation is that the different response between echolocation and communication vocalizations was a result of a different response to physiological stress but not to acoustic masking. As mentioned above, the adjustments in echolocation vocalizations may be a stressor response to traffic noise. Animals may improve their vigilance in stressful conditions (Rozan et al., 2008). Bats use echolocation calls mainly for environmental perception and use communication calls for social communication. It is more likely that bats change their echolocation calls than communication calls to increase vigilance. Alternatively, the bats under stress may simply emit more echolocation calls without any function.

### Implications for Conservation

The affect of anthropogenic noise on bat fitness remains unclear. Traffic and/or gas compressor station noise reduced the foraging activity and efficiency in *Antrozous pallidus* (Bunkley and Barber, 2015), *Myotis daubentonii* (Luo et al., 2015b), *M. myotis* (Schaub et al., 2008; Siemers and Schaub, 2011), or *Tadarida brasiliensis* (Bunkley and Barber, 2015), but not that of *M. californicus, M. cillolabrum, M. lucifugus,* or *Parastrellus Hesperus* (Bunkley and Barber, 2015). The impacts of noise appears to vary with species and the behavioral/physiological state of the animal (Luo et al., 2014). We found that traffic noise increased the echolocation activity of *V. sinensis* nearly twofold. Echolocation is energetically costly for resting bats and increasing echolocation vocalization could result in higher energy costs (Speakman et al., 1989). We argue that even if noise exposure may not decrease fitness related to foraging, it may increase the energy expenditure due to more vocalization in bats. If the increased vocal activity reflects a stress response the long-term stress could increase incidence of disease (Romero and Butler, 2007).

In conclusion, this study shows that the Asian particolored bats change the temporal features of echolocation vocalizations but not those of communication vocalizations in response to traffic noise. These findings suggest that vocal responses to anthropogenic noise can be inconsistent among different types of vocalizations and that the degree of spectral overlap between animal vocalizations and noise does not necessarily predict the level of vocal response to noise. Future work will advance these results by performing behavioral and physiological experiments to examine why bats increase their echolocation activity when they are exposed to noise.

## Acknowledgements

We thank Heng Liu for bat collection and Bailu Si for help with data analysis.

## Author contributions

S.S., A.L., T.J. and J.F. conceived and designed the experiment; S.S., and, X.Z. performed the experiment, S.S. analyzed the data; S.S. and A.L. drafted the paper; W.M. and J.F. revised the paper.

---

**Figure S1.**
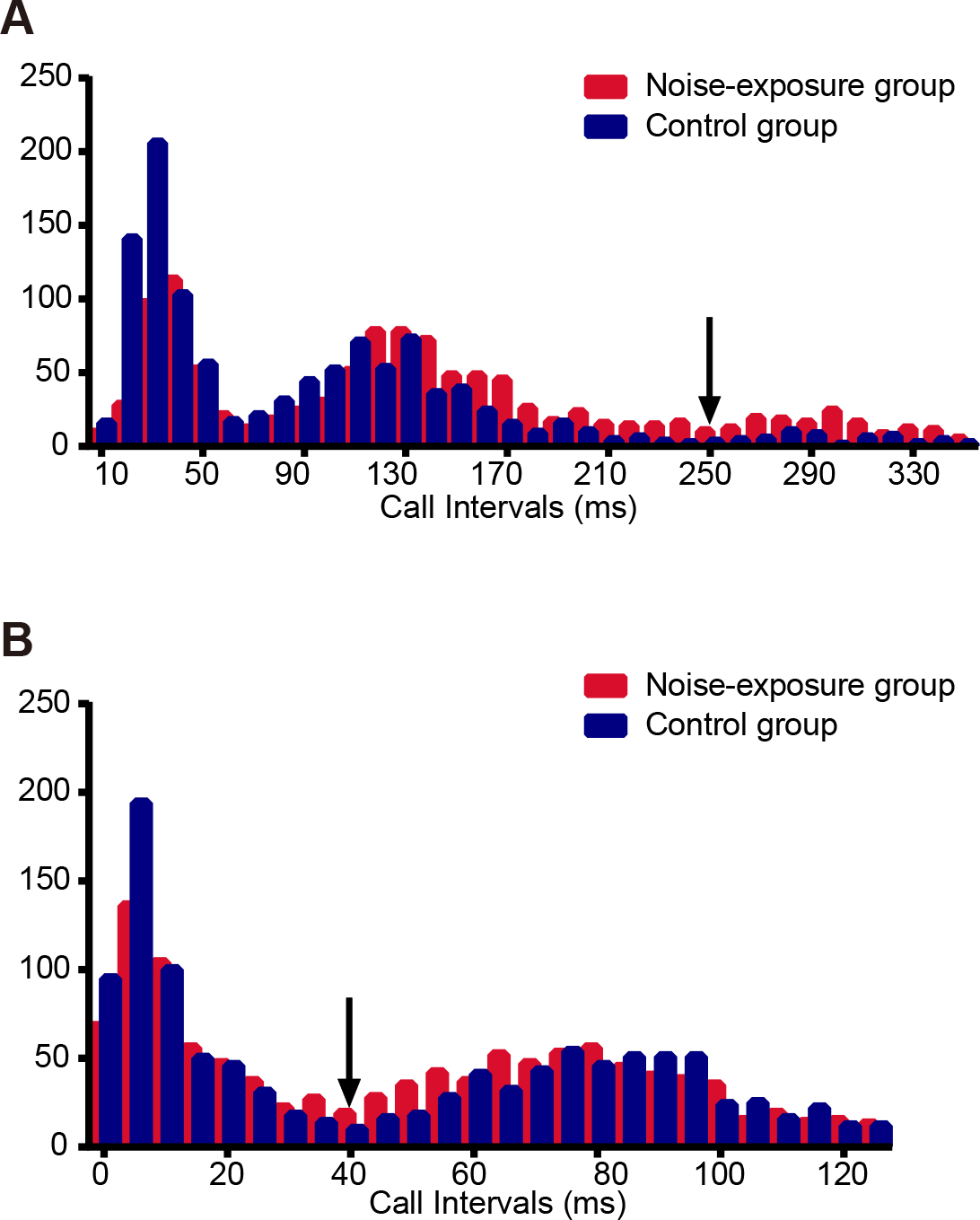
Frequency distribution of inter-call intervals for echolocation. (A) and communication (B) call sequences in *Vespertilio sinensis.* The arrow indicates the boundary of call sequence.

**Table S1.**
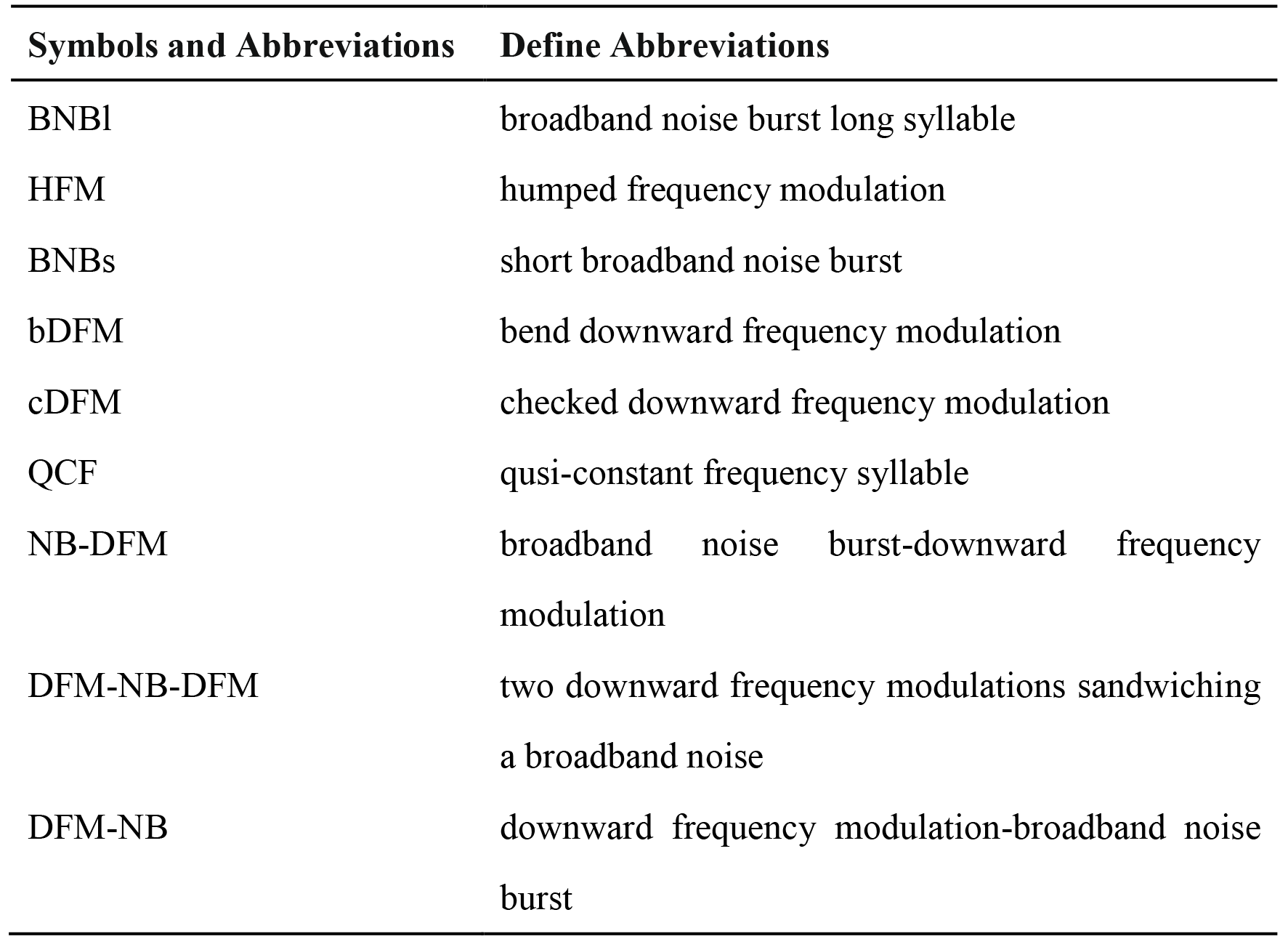
List of Symbols and abbreviations of communication syllables of *Vespertilio sinensis*.

